# A scalable implementation of the recursive least-squares algorithm for training spiking neural networks

**DOI:** 10.1101/2022.09.26.509578

**Authors:** Benjamin J. Arthur, Christopher M. Kim, Susu Chen, Stephan Preibisch, Ran Darshan

## Abstract

Training spiking recurrent neural networks on neuronal recordings or behavioral tasks has become a popular way to study computations performed by the nervous system. As the size and complexity of neural recordings increase, there is a need for efficient algorithms that can train models in a short period of time using minimal resources. We present optimized CPU and GPU implementations of the recursive least-squares algorithm in spiking neural networks. The GPU implementation can train networks of one million neurons, with 100 million plastic synapses and a billion static synapses, about 1000 times faster than an unoptimized reference CPU implementation. We demonstrate the code’s utility by training a network, in less than an hour, to reproduce the activity of > 66, 000 recorded neurons of a mouse performing a decision-making task. The fast implementation enables a more interactive *in-silico* study of the dynamics and connectivity underlying multi-area computations. It also admits the possibility to train models as *in-vivo* experiments are being conducted, thus closing the loop between modeling and experiments.

## 1 Introduction

Cognitive functions involve networks of interconnected neurons with complex dynamics that are distributed over multiple brain areas. One of the fundamental missions of system neuroscience is to understand how complex interactions between large numbers of neurons underlie the basic processes of cognition.

An increasingly popular data-driven modeling approach for investigating the neural mechanisms that support behavioral tasks is to train neurons in an artificial neural network to reproduce the activity of recorded neurons in behaving animals [1, 2, 3, 4, 5, 6]. Such network models can range from purely artificial networks that are far from being biological [2, 3, 4, 7], to biophysical neuronal networks that include spiking activity [8] of different cell types that operate in a brain-like dynamical state [9]. The neural dynamics and the connectivity structure of the trained network can then be analyzed to gain insights into the underlying neural mechanisms. For instance, [2] demonstrated that memory-related sequential activity can be produced in highly heterogeneous but partially structured recurrent neural networks, [4] showed that decision-related information can be gated from distracting inputs by forming increasingly stable fixed points that move away from the decision boundary, and [9] provided general circuit mechanisms for spreading task-related neural activities from a small subset of trained neurons to the rest of neurons in the network.

With the increase in the size of experimentally recorded neural data sets, the ability to fit the activity of neurons in model networks is becoming a challenge. For example, the number of simultaneously recorded neurons in behaving animals has been increasing in the last few years at an exponential rate [10]. At present, it is possible to simultaneously record in a single session about 1,000 neurons using electrophysiology, and up to 100,000 using calcium imaging in behaving animals [11]. When combining several sessions of recordings, the amount of data becomes huge and will likely grow to millions of recorded neurons in the near future.

Here, we developed a scalable implementation of the recursive least-squares algorithm (RLS) to train spiking neural networks of tens to hundreds of thousands of neurons to reproduce the activity of neural data. RLS [12] was initially applied to train the outputs of a recurrent neural network for performing complex tasks, such as implementing 3-bit memory or generating motor movements (also known as FORCE [7]). Subsequently, RLS was adopted for training the individual neurons within a recurrent neural network to reproduce target neural activities. Examples of target activities include activity of neurons recorded from the brain [2, 4, 3], chaotic activity of a random network [13], teacher networks [14] and arbitrary functions [8]. Although most existing studies apply RLS to rate-based networks, it can also be implemented in spiking neural networks for performing complex tasks [15] and reproducing neural activities [8, 16, 9].

Starting with the code in [9], we generalized it to include different models of single cell dynamics, as well as extended it to account for external noise. We then optimized it for run-time speed. Finally, we demonstrated its performance by training more than 66,000 spiking neurons to reproduce the activity of recordings from multiple brain areas of mice using Neuropixels probes [17] in a decision making task [18, 19]. Fitting these data, which were sampled at 20ms for 3 seconds, takes about 10 hours on a multi-core CPU and less than an hour on a GPU. The code is freely available.

## 2 Results

We implemented the RLS algorithm to train individual neurons within a large spiking recurrent neural network to reproduce a pre-determined target activity (Figure 1A). Specifically, we considered networks of integrate-and-fire spiking neurons connected by synapses with varying spike-filtering time scales but no explicit spike-propagation delays, in which the neurons could fire irregularly due either to recurrent interactions, known as the fluctuation-driven regime [20, 21, 22], or to external noise, or both. We chose the standard leaky integrate-and-fire neuron as our model neuron. In the code, the cell model is a plugin, making it easy for users to customize, and we provide code that implements all five of the Generalized Leak Integrate and Fire (GLIF) models described in [23].

**Figure 1:**
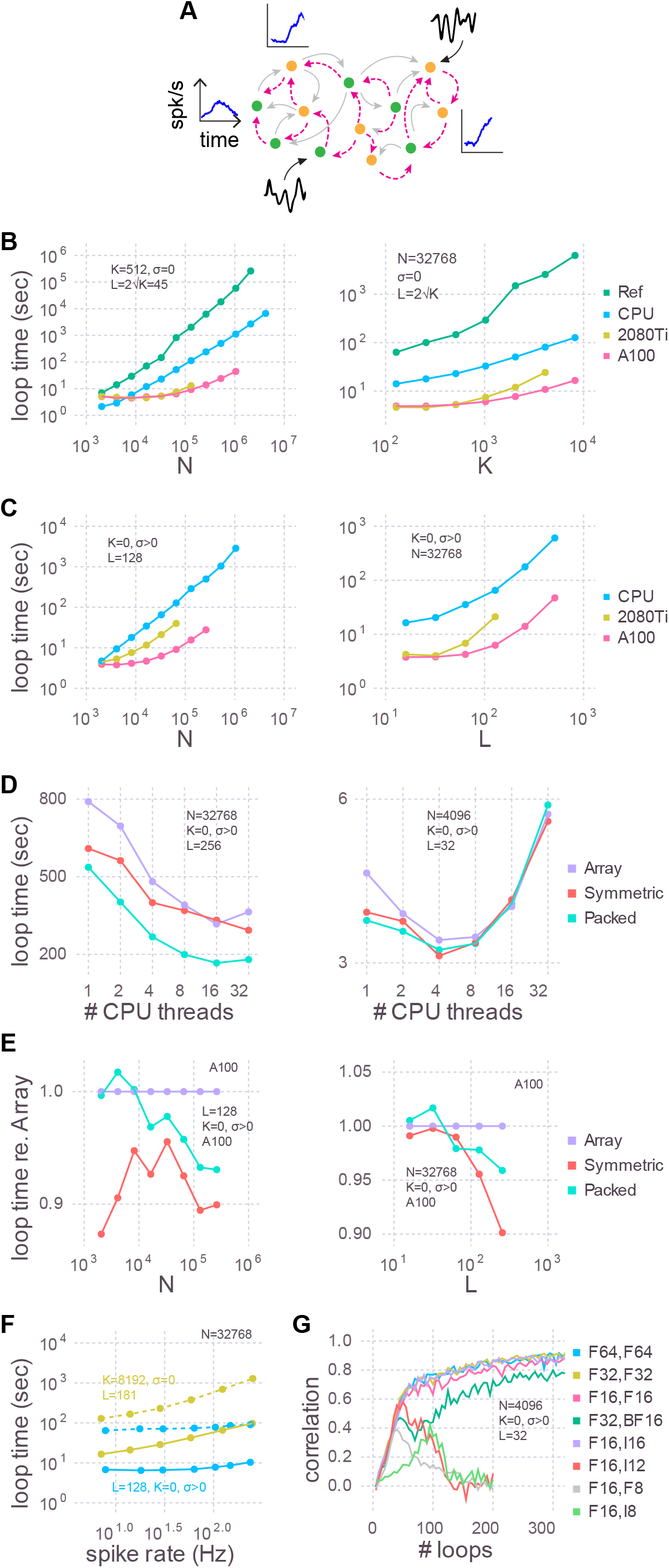
Training speed for models of various sizes. **(A)** Cartoon of a recurrent spiking neural network. Excitatory (green circles) and inhibitory (yellow circles) neurons have plastic (magenta lines) and, optionally, static (grey lines) connections to each other. Each neuron receives, on average, *K* static and *L* plastic connections. Associated with each neuron are the target activity patterns (blue insets; only three shown). For models with no static connections (*K* = 0) we injected into each neuron a white noise with variance *σ* (black insets; only two shown). **(B)** The time taken by each training iteration for models with static connections versus the number of neurons (*N* ; left) or the number of static connections (*K*; right). Compared are the single-threaded CPU code used by [9] (Ref), our optimization of the same CPU code (CPU), and our refactoring of the same algorithm for GPUs using a consumer grade card (2080Ti) and an enterprise grade board (A100). In each case we tested the model sizes up to the largest that would fit in memory: 768 GB for the CPU, 11 GB for the 2080Ti, and 80 GB for the A100. The reference code used 64-bit floating point numbers and a full dense array for the large ***P*** matrix; all new code presented here uses 32-bit floating point numbers and a packed symmetric array for ***P***. **(C)** Similar to **(B)** but with a Gaussian noise model instead of static connections (*K* = 0, *σ >* 0), when varying the number of neurons (left) or the number of plastic connections (*L*; right). **(D)** Strong scaling of the optimized CPU code for large (left) and small (right) models. Purple: code stores the ***P*** matrix in a full dense matrix. Red: symmetric matrix. Cyan: packed symmetric matrix. **(E)** The effect of the matrix storage format on the GPU code as a function of the number of neurons (left) or plastic synapses (right). **(F)** The time taken for each training iteration as a function of spike rate. The latter was varied with the external input to neurons (*X*_*bal*_ in Eq. (7)). Solid lines indicate GPU code and dotted are the CPU code. **(G)** The effect of storage precision on learning. The first column in the legend indicates the bit precision of all state variables *except* ***P***, which is indicated in the second column. For integer types, ***P*** was scaled by 2^*nbits*−2^.

The learning objective was to train the synaptic current *u*_*i*_(*t*) of each neuron *i* = 1, …, *N* such that it followed a target activity pattern *f*_*i*_(*t*) on a time interval *t* ∈ [0, *T*] (see Appendix: Recursive Least Squares). These activity patterns were extracted from the peri-stimulus time histograms (PSTHs) of recorded neurons in the brain (see Appendix: Generating target trajectories). To trigger a target response, each neuron in the network was stimulated by a constant input with random amplitude for a short duration. These external stimuli were applied to all neurons simultaneously, such that the network was set to a specific state at the end of stimulus duration. The synaptic weights were trained with this stimulated network state as the initial state, which allowed the trained network to produce the target response whenever the stimulus was applied to reset the network state. We treated every neuron’s synaptic current as a read-out, which made our task equivalent to training *N* recurrently connected read-outs. For the current-based synapses considered in this study, neuron *i*’s synaptic current *u*_*i*_ can be expressed in terms of the spiking activities of other neurons *r*_*j*_, *j* = 1, …, *N* through the exact relationship *u*_*i*_ = ∑_*j*_ *W*_*ij*_*r*_*j*_ (see Eqs. (9) and (11) for details). Therefore, we adjusted the incoming synaptic connections *W*_*ij*_, *j* = 1, …, *N* to neuron *i* by the RLS algorithm in order to generate the target activity. Specifically, we randomly chose *L* plastic connections per neuron and trained only these connections. This training scheme allowed us to set up independent objective functions for each neuron and update them in parallel (see Appendix: Recursive least-squares). The linearity of *u* in terms of *r* makes it possible to directly apply the RLS algorithm to train the synaptic activity. Alternatively, the firing rates of neurons can also be trained if an appropriate F-I curve (Eq. (1)) for the neuron model is used in the RLS algorithm. This procedure, however, involves differentiating the nonlinear F-I curve, which can slow down learning during subthreshold activity, as shown previously [8].

Besides the plastic connections, each neuron can also receive an average of *K* pre-synaptic static inputs. These connections are referred to as ‘static’ because they remain unchanged during the learning process. The average static synaptic weights were chosen such that the network was in the balanced state (see Appendix and [9] for further details). In addition, an external noise was optionally injected to each neuron, with a variance of *σ*^2^.

In the case where the neurons are driven by external noise and there are no static connections (*K* = 0, *σ* > 0), the number of plastic weights (*L*) is only bounded by *N* (and memory). However, in the presence of static weights, we follow our previous work [9] and choose them such that 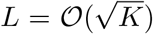. As discussed in [9], this relationship, along with the scaling of 1*/L* for each plastic synapse, which is comparable to the static synapses (i.e. 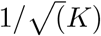), facilitates learning without disturbing the balanced state of the network.

Starting with a working implementation of the algorithm [9], we profiled the code to find slow sections, optimized those lines for performance using the strategies below, and iterated until there were no further easy gains to be had. We achieved almost two orders of magnitude improvement in run times for large networks using the CPU alone compared to the reference code (Fig. 1B), and another order of magnitude or two by refactoring to use a GPU. These trends held true when random static connections were replaced with a Gaussian noise model (Fig. 1C). The advantage of this noise model is that run times are relatively independent of firing rates (Fig. 1F).

### 2.1 Optimization strategies

We used the Julia programming language [24] since rapid prototyping and fine-grained performance optimizations, including custom GPU kernels, can be done in the same language. Several strategies and techniques were used to make the code performant, both in terms of run time as well as memory use. Benchmarking was performed on synthetic data consisting of sinusoidal target functions with random phases.

### 2.2.1 Parallel updates of the state variables

Pre-synaptic weights and neuronal voltages can be updated in parallel as they are all independent of each other. Synaptic currents can also be updated in parallel, except during an action potential, when different threads might overwrite each others updates to a post-synaptic neuron’s current if multiple of its pre-synaptic neurons spike simultaneously. In the CPU code, we used multiple threads to loop over the neurons to update the weights, voltages, and the exponential decay of currents. As Julia does not support atomic operations on elements of a vector, and locking mechanisms can be slow, a subsequent non-threaded loop was used to update the post-synaptic currents to avoid the race condition when a spike occurred.

We benchmarked CPU multi-threading on a machine with 48 physical cores and found that performance plateaued after 16 threads for a large model (Figure 1D). For small network models there was an optimal number of threads, and using more threads was actually slower. As there is no communication between threads, the optimum number is presumably determined by the balance between the overhead in launching each thread and the time spent performing computations therein. There is also a dependence of loop times on spike rate, due to the second non-threaded loop, the slope of which is proportional to *K* + *L* (Figure 1F).

Given that GPUs are purpose-built to thread well, we investigated whether the RLS algorithm would scale better with them. Since accessing individual elements in a vector is very slow on a GPU, due to each kernel call incurring a large overhead, we made a functionally equivalent copy of the CPU code and refactored it to use vector operations instead of for-loops over elements. Specifically, to reset the membrane potential in a performant way when a spike occurs, we replaced the for-loops in the CPU code, and the if-statements therein, with a broadcasted ifelse statement that inputs a boolean vector indicating which neurons spiked. Doing so results in the membrane potential of all of the neurons which do not spike being “reset” to its current value, but this is still faster even for low spike rates. To update the post-synaptic neurons, we moved the for-loop to be inside a custom GPU kernel where CUDA atomic instructions are available to handle the race condition. Just one kernel call is hence made, thereby minimizing the overhead. A dependency of loop time on spike rate also exists, just like in the CPU code, due to the atomic locking.

Whereas the CPU run times were linear with model size, GPU performance was flat below a certain size. This is likely because models that are sufficiently small don’t use all the parallelism that the GPU provides. The NVIDIA A100, for example, has 108 multiprocessors and each one can execute 32 threads simultaneously, yielding a total of 3456 threads.

#### 2.1.2 Symmetric and packed arrays

The RLS algorithm uses the running estimate of the inverse of the correlation matrix of the activity of the pre-synaptic neurons (plus regularization terms), which for each of the *N* neurons is a symmetric matrix of size *L* × *L* that we denote as ***P*** (see Eq (18)). To perform mathematical operations on ***P***, as well as other state variables, we utilized Basic Linear Algebra Subprograms (BLAS), a highly-engineered library of mathematical operations commonly used in high-performance computing. While some BLAS routines specialized to operate on symmetric matrices are faster, others are slower. Consider the function syr, for example, which computes ***A*** = *α****xx***^*T*^ + ***A***, where ***A*** is a symmetric matrix, ***x*** is a vector, and *α* is a scalar. Here ***A*** is being updated, and since it is symmetric, there are only half as many elements to update compared to ger which computes the non-symmetric counterpart. syr is hence typically faster than ger. Conversely, symv computes ***y*** = *α****Ax*** + *β****y***, where ***A*** is a symmetric matrix, ***x*** and ***y*** are vectors and *α* and *β* are scalars. Here every element of ***A*** must be accessed to update ***y***. Since extra logic must be used to ensure that indexing operations only access a particular triangle, symv is typically slower than gemv. We found that on balance it was slightly faster to use routines which operate on the symmetric ***P*** matrices (Figure 1D,E), particularly for models with large number of plastic synapses.

Further, ***P*** consumes by far more memory than any other variable since its footprint scales as *N* × *L*^2^. All other state variables are only one or two dimensional. Packing the columns of just the upper or lower triangle by concatenating them into a vector saves close to half the memory, thereby permitting models to be proportionately larger. Though a bit slower on the GPU overall compared to their unpacked counterparts (symv and syr; Figure 1E), BLAS routines specialized for packed symmetric matrices (spmv and spr) are much faster on the CPU (Figure 1D) for large models. We speculate that this performance difference is due to the sophisticated hierarchical caches on a CPU being better utilized with packed matrices, compared to a GPU.

#### 2.1.3 BLAS, pre-allocated memory, and pre-computed division

We found that simple refactorings of our CPU code to directly use BLAS resulted in substantial performance gains. For example, the RLS algorithm computes ***k*** = ***P r***, which is a product of the symmetric matrix, ***P***, and the presynaptically filtered spike trains, ***r*** (see Eqs (9) and (18)). Preallocating and reusing ***k*** and then calling *mul*!(***k, P***, ***r***), which is a thin wrapper around BLAS’ gemv matrix-vector multiplication function, is faster and uses less memory than doing the dot product directly.

A further performance improvement was realized by using Intel’s Math Kernel Library (MKL) for the CPU, which is a superset of BLAS hand-crafted for the x86 architecture, instead of the default cross-platform OpenBLAS. Decrements in loop times were most pronounced for models with *K >* 0. Specifically, for a model with *N* = 32768, *K* = 1638, *L* = 81 we saw a 59% reduction using MKL, whereas for *N* = 32768, *K* = 0, *L* = 128 there was only a 5.8% improvement, about 10-fold less.

BLAS functions frequently input multiplicative constants, forcing the user to manually do a division ahead of time if the constant is in the denominator. Following this lead for the sections of code that do not use BLAS directly, we precomputed, just once, the inverse of the synaptic currents and membrane voltage time constants as they are in the denominator of the equations that govern the neural dynamics. Loop times were about 2% quicker performing this multiplication, instead of the corresponding division, for the CPU code.

For the GPU version of our code we wrote our own GPU kernels which batched several BLAS routines, specifically gemv and ger plus their symmetric (symv, syr) and packed symmetric equivalents (spmv, spr). Such batched GPU kernels are critical to our learning algorithm since we apply the RLS algorithm to each of the N trained neurons, and calling a non-batched kernel N times would incur a huge performance penalty due to the overhead of calling functions on GPUs. The solution was to write new BLAS kernels that internally iterate over N, which was necessary as NVIDIA only provides batched implementations of gemm, gemv, and trsm.

#### 2.1.4 Reduced precision number formats

Our original reference code in [9] used 64-bit floating point precision for all variables, which can represent numbers up to 1.8 × 10^308^ with a machine epsilon of 2.2 × 10^−16^ around 1.0. We found that the correlations between target activity patterns and learned synaptic currents, which measure the accuracy of the training, are just as high and require the same number of iterations using 32-bit floats, whose range is only 3.4 × 10^38^ and machine epsilon is 1.2 × 10^−7^ (Figure 1G). Doing so not only permits models twice as large to be trained, but also yields quicker loop times on CPUs and consumer grade GPUs (data not shown).

Our custom batched BLAS kernels for the GPU can operate on floating point numbers of any precision, and even integers, unlike the CPU BLAS libraries. To further reduce memory consumption, and hence increase model size, we tried Float16, whose range is only 6.5 × 10^4^ and machine epsilon is 9.7 × 10^−4^, and found that models can be trained almost as accurately as with 64 and 32 bits. To see if the small decrement in correlation could be recovered with a different partitioning of the bits between the exponent and the fraction, we tried BFloat16, which uses eight bits for the exponent, just like Float32, and hence has the same range, instead of Float16’s five. Correlation coefficients for BFloat16 were worse. As there are no hardware-supported 16-bit floating point formats that allocate more bits to the fraction, we tried scaling ***P*** by 2^14^ and storing it in a 16-bit integer, effectively giving us 14 fractional bits. There is no overflow with this scheme, as ***P*** typically ranges from -1 to 2. Scaled 16-bit integers had correlations that were indistinguishable from Float32 and Float64. Furthermore, there is no speed penalty with 16-bit integers.

Modern hardware does not support 8-bit floats (though see the forthcoming NVIDIA H100 GPU), but they can be simulated in software with, for example, MicroFloatingPoint.jl. We tried 8-bit floats with two, three, four, or five exponential bits and found that correlations were best with four and reached a peak around half that of 32 and 64 bits and then declined. Scaling ***P*** by 2^6^ and storing it in an Int8 resulted in a correlation peak of similar height. While Int8 required more iterations to reach the peak than Float8, the wall clock time was actually less as the loop times were much shorter due to native hardware support.

There is also no direct hardware support for 12-bits, but given the large difference between our 8 and 16 bit results, we simulated them in software by using an Int16 [sic] scaled by 2^10^. Since the magnitude of ***P*** is less than two, the four most significant bits here are entirely unused, and the 10 least significant are used as the fraction. As with scaled Int8, we observed a peak in the correlation coefficient, this time a bit larger at about two-thirds that of 32 and 64 bits, followed by decline. A 12-bit integer is not an unreasonable type to imagine using to reduce memory consumption, as two Int12 will pack into three bytes. In fact, UInt12Arrays.jl is a Julia package which does precisely this for unsigned 12-bit integers.

In summary, we recommend using Float32 unless the model does not fit in memory, in which case Int16 can be used. If it still does not fit and a drop in correlation can be tolerated, we recommend using Int8, or, time permitting, developing a proper Int12 package.

### 2.2 Application

With a fast RLS codebase in hand, we next demonstrated that large models can be successfully trained to recapitulate the dynamics in real-world big data sets. We used 66,002 peri-stimulus time histograms (PSTHs) of neurons, recorded using Neuropixels probes [17] from multiple brain areas of mice performing a delayed-response task (Figure 2A; [19, 25]). We first converted the PSTHs to the corresponding underlying synaptic currents by inverting the activation function of leaky integrate-and-fire neurons in the presence of noise (see Appendix: Generating neural trajectories). The synaptic currents were then used as the target functions, and external noise was used instead of static recurrent connections (*K* = 0, *σ* > 0; see Appendix: Network dynamics). As our goal was to show the scalability of the code and not to study the trained network, we did not use any prior knowledge on mesoscopic connectivity between or within brain regions, but simply initialized the plastic weights by randomly connecting neurons in an Erdós–Rényi graph. In addition, the trained plastic weights in our network were allowed to flip signs, and hence possibly violate Dale’s law. However, in principle one could use mesoscopic connectivity to constrain the plastic weights and the RLS algorithm could be further developed to obey Dale’s law [26].

**Figure 2:**
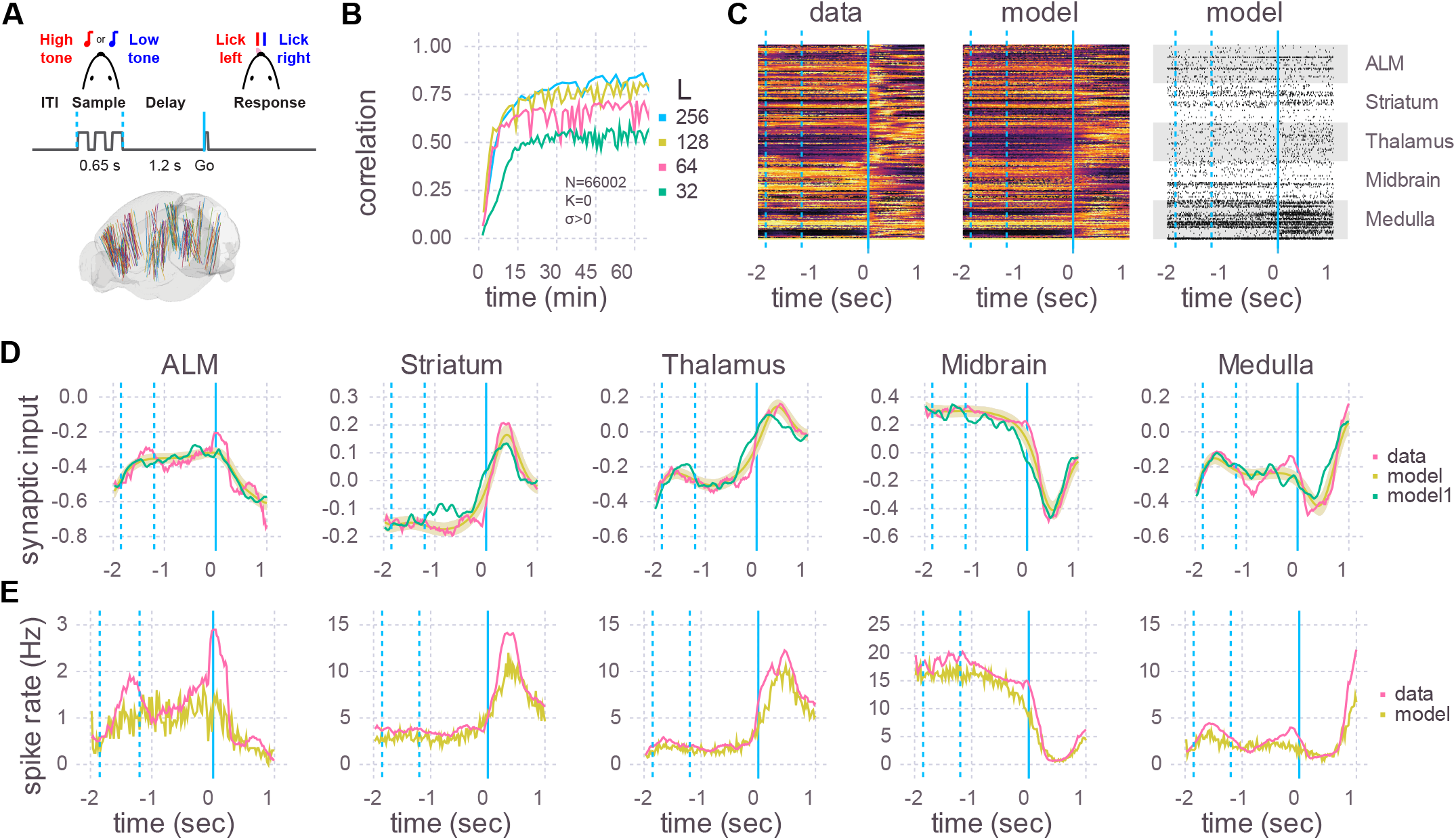
Application to Neuropixels data. **(A)** Top: Schematic of experimental setup. Mice learned to lick to one of two directions (left/right) after a delay period, depending on which of two tones were played. Bottom: Multiple Neuropixels probes, each with hundreds of recording sites, were placed in the brain while a mouse was performing the task. [19] **(B)** A recurrent spiking neural network with 66,002 neurons and no static connections (*K* = 0, *σ >* 0) was trained to learn the neural activity patterns. Initial learning rate and final plateau level both increased with the number of plastic connections (L). The number of training iterations was 375, 330, 235, and 110, respectively, for L = 32, 64, 128, and 256. **(C)** Heat maps of the peri-stimulus time histograms (PSTHs) of the activity patterns in the data (left) and the trained network from B with *L* = 256 (middle) and one realization of spike trains of the same trained network (right). 50 neurons are shown for each of five brain areas. Vertical blue lines are the same as in **A**, where a time of 0 sec corresponds to the Go cue. **(D)** Learned synaptic currents averaged over 1000 trials for one exemplar neuron from each of the five brain areas. Vertical blue lines are the same as in **A**, and the time axis is the same as in **C**. A single realization (non-averaged over trials) is also shown (model1). **(E)** Same as D but for PSTHs.

We then used the RLS algorithm to train the neurons in the spiking network to follow the target functions. The correlation between the learned and target synaptic currents increased with the number of plastic inputs (*L*), and reached a plateau in half an hour (Figure 2B). 256 plastic synapses was the largest that would fit in the 80 GB of memory in an A100 GPU with N=66,002. For comparison, a model of that size would take multiple days on a single thread of a CPU using the reference code of [9]. After training, we ran the spiking network multiple times, with the plastic weights kept frozen to the trained values (i.e., the weights are no longer changed with the RLS algorithm), and compared the learned synaptic currents (Figure 2D) and PSTHs (Figure 2C,E). The activity of neurons in the trained network showed a close correspondence with the activity patterns of the recorded neurons.

Note that in this experiment each trained neuron successfully learned two activity patterns corresponding to lick right and lick left trials. Learning more than two patterns would likely require additional plastic synapses, which could potentially limit the task complexity that can be achieved using a single GPU.

## 3 Discussion

We present optimized CPU and GPU implementations of the recursive least-square (RLS) algorithm for training spiking neural networks. Our code can simulate and train a spiking network consisting of about one million neurons and 100 million synapses on a single modern high-end workstation.

Our updated code is significantly faster than the previous version [9], as demonstrated by the green line in Fig. 1B. Networks consisting of millions of neurons can be trained 1000 times faster using the new code. We benchmarked the code for various scaling numbers (N, K, L) and also expanded its capabilities to include external noise (*σ*), allowing for training of both balanced [20, 27] and generic spiking neural networks. This increased efficiency means that large networks of a million neurons can now be trained in a matter of hours instead of weeks, thereby greatly speeding up the scientific discovery process and more efficiently using resources. Additionally, fast training times enables models to be created on the fly, allowing for *in-silico* training while *in-vivo* experiments are being conducted, thus closing the loop between modeling and experiments. For example, predictions could be made about perturbation experiments by fitting models to data and suggesting perturbation protocols in real-time.

While we used simple integrate-and-fire neurons, our code can be easily extended to include more realistic neurons using a cell-model plugin system that provides users the means to provide custom code. A simple integrate-and-fire neuron allowed us to easily convert PSTHs to synaptic currents using the known F-I curve (Eq.(1); [28]). It remains as future work to develop principled methods to convert the PSTHs to synaptic currents in more complex neuron models that include slow dynamic variables, such as adaptation currents or time-dependent spike thresholds [23]. However, the RLS algorithm used here allows one to train the synaptic currents of GLIF neurons, if the target synaptic currents are already available (see [8] for training a network of Izhikevich neurons).

Additionally, although the plastic weights in our networks were random, our framework does not exclude adding complexity to the network architecture, such as including layered cortical networks or even different cell types, or using known mesoscopic connectivity when initializing the plastic weights in the network. This flexibility is again achieved through a set of plugins through which the user can provide custom code to define the adjacency matrices.

The RLS algorithm is designed to minimize the discrepancy between the PSTHs generated by the model neurons and the recorded neurons. It does not optimize for other spiking statistics, such as the coefficients of variation for interspike-intervals or Fano factors. These statistics can be influenced by adjusting the network’s hyper-parameters, such as the level of noise applied to each neuron (*σ*). Tools are available to assess the network’s performance against these other measures while varying the hyper-parameters (e.g. https://github.com/INM-6/NetworkUnit).

The size of the networks is limited by memory usage, which mainly depends on the size of the matrix ***P*** used in the RLS algorithm (see Eq (18)). This matrix scales linearly with the number of neurons and quadratically with the number of plastic synapses (*N ×L*^2^). We found that about *L* ≈ 100 synapses per neuron suffices to train the network to reproduce the activity of neurons recorded in mice performing a delayed response task (Figure 2B). However, the number of plastic synapses needed to train the network is expected to increase with the number of tasks to be learned, as well as the complexity of neuronal dynamics in each task. Therefore, how large the network model would have to be could depend on the number and the complexity of the tasks to be learned.

Advances in GPU computing and strong interest in neuromorphic computing have led to various efficient implementations of spiking neural networks. Recent work that implements simulations of spiking neural networks in GPUs include the following: a code-generation based system that generates CUDA code for GPU (GeNN [29]) and a popular Python-based simulator for spiking neural networks, Brian2, extended for generating CUDA code directly (Brian2CUDA [30]) or through GeNN (Brian2GeNN [31]). Similarly, highly efficient CPU-based simulations of spiking neural networks can be implemented in NEST (NEural Simulation Tool [32]). Spiking neural networks can also be implemented in neuromorphic hardware as demonstrated in SpiNNaker [33], Intel’s Loihi [34] and TrueNorth [35]. See [36] and references therein for benchmarks for these systems. We note that, although the aforementioned systems allow for biologically plausible plasticity [31] and for learning complex tasks [34, 35], the contribution of our work is different in that we developed an efficient algorithm for learning to generate activity patterns in recurrently connected spiking neural networks.

Finally, our code implementation was tailored to be used on a single computer, instead of on multiple computers, such as in [37]. This enabled fast execution speed, thanks to the absence of interprocess communication overhead, but limited the network size due to memory limitations. Specifically, the CPU implementation uses threads (not processes), which have precisely zero communication overhead because they share memory. The GPU implementation has currently only been tested on a single GPU. However, modern hardware and the associated software toolkits support an abstraction of a unified memory across multiple GPUs within a single computer that is backed by high-speed interconnects. If in future, the size of GPU memory does not increase at a rate comparable with advances in neural recording technology, we plan to investigate how much performance is decremented if our code is refactored to use multiple GPUs within the same workstation.

## Data availability statement

The main code repository is located at https://github.com/SpikingNetwork/TrainSpikingNet.jl.

We also release two new Julia packages that it depends on, and put them in separate repositories for easy composability with other code bases: one for packing symmetric matrices and the other for batching (packed) symmetric BLAS routines on the GPU. See https://github.com/JaneliaSciComp/SymmetricFormats.jl and https://github.com/JaneliaSciComp/BatchedBLAS.jl, respectively.

For NeuroPixels physiological data see [19]. The corresponding target synaptic currents plus the parameters file that we used for these data to generate Figure 2 are at https://github.com/SpikingNetwork/TrainSpikingNet.jl/releases/download/v0.1.0/figure2-neuropixels.tar.xz.

### Author contributions

CMK and RD conceived of the algorithm, CMK provided and helped understand the reference implementation, BJA optimized the code for performance, with the assistance of CMK and RD. SC did the experiments and collected and analyzed the Neuropixels data, BJA prepared the figures, BJA, CMK, SP, and RD wrote the manuscript.

## Acknowledgements

We would like to thank Shaul Druckmann, Nuo Li, Yi Liu and Karel Svoboda for sharing the Neuropixels data set.

## 4 Appendix

Here we provide technical details on how spiking neural networks were trained. A large part of following presentation was adopted from the previous paper of the authors [9].

### 4.1 Generating target neural trajectories from PSTHs

For each recorded neuron, we converted its PSTH to target pre-synaptic activities to be used for training the synaptic inputs. Specifically, in Figure 2 we obtained for each spike rate *r*_*it*_ (*i* = 1, …, *N*, and *t* = *T*_*init*_ + ≈ *t*, …, *T*_*end*_ where *T*_*init*_ = −2 and *T*_*end*_ = 1) the mean synaptic input *f*_*it*_ that needs to be applied to the the leaky integrate-and-fire neuron to generate the desired spike rate. To this end, we numerically solved the nonlinear rate equation

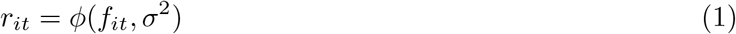

where

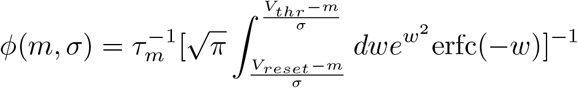

is the transfer function of the leaky integrate-and-fire neuron given mean input, *m*, and variance of the input, *σ*^2^ [38, 28]. This conversion yielded a set of vectors of target synaptic inputs **f**_1_, ∦, **f**_*N*_ ∈ R^*T*^ where *T* = (*T*_*end*_ − *T*_*init*_)*/*≈ *t* for the recorded neurons. Note that in Figure 2 we did not simulate the recurrent static connections (*K* = 0). Simulating both static connections and the external noise would require estimating *σ* in Eq.(1). This can be done by assuming Poisson firing statistics and calculating *σ* given the average connectivity and rates of the populations in the network [28]. Alternatively, it is possible to directly estimate *σ* from simulating an initial network that fits the average rates and level of noise (e.g. the Fano factor) observed in the data [9]. The results presented in this paper are done using the latter approach.

### 4.2 Network connectivity

In Figure 1A, the spiking neural network consisted of randomly connected *N*_*E*_ excitatory and *N*_*I*_ inhibitory neurons. They were recurrently connected as in [9]. In short, the recurrent synapses consisted of static weights **J** that remained constant throughout training and plastic weights **W** that were modified by the training algorithm. The static synapses connected neuron *j* in population *β* to neuron *i* in population *α* with probability *p*_*αβ*_ = *K*_*αβ*_*/N*_*β*_ and synaptic weight 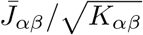, where *K*_*αβ*_ is the average number of static connections from population *β* to *α*:

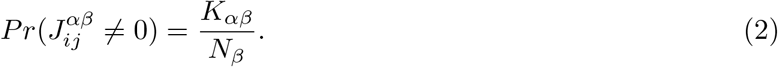

The strength of plastic synapses, 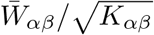, was of the same order as the static weights. However, the plastic synapses connected neurons with a smaller probability:

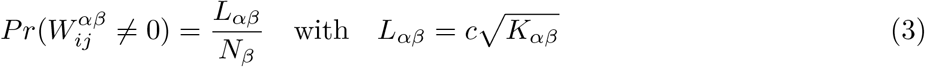

which made the plastic synapses much sparser than the static synapses [39, 9]. Here, *c* is an order 1 parameter that depends on training setup.

The static and plastic connections were non-overlapping in that any two neurons in the network can have only one type of synapse.

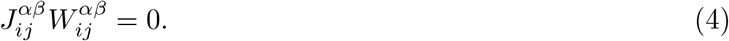

Keeping them disjoint allowed us to maintain the initial network dynamics generated by the static synapses and, subsequently, introduce trained activity to the initial dynamics by modifying the plastic synapses.

The static recurrent synapses were strong in that the coupling strength between two connected neurons scaled as 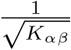, while the average number of synaptic inputs scaled as *K*_*αβ*_. As a result of this strong scaling, the excitatory and inhibitory synaptic inputs to a neuron from static synapses increased as *K*_*αβ*_, and thus were much larger than the spike-threshold for a large 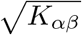. However, the excitatory and inhibitory currents were dynamically canceled, and, together with the external input, the sum was balanced to be around the spike-threshold [20, 9].

In contrast to the static synapses, each trained neuron received only about 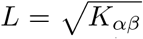 plastic synapses. This made the plastic synapses much spa≈ rser than the sparse static EI connectivity (e.g., with *K* = 1000 static synapses, there are of the order of 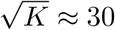 plastic synapses per neuron). Consequently, the EI plastic inputs of the initial network were independent of *K*_*αβ*_ and substantially weaker than the EI balanced inputs for a large *K*_*αβ*_. After training the plastic synapses, the total synaptic input, i.e., the sum of plastic and balanced inputs, to each trained neuron was able to follow the target patterns. With this scaling of plastic synapses, training was robust to variations in the network size, *N*, and the number of pre-synaptic connections, *K*_*αβ*_ (Fig. 1B).

In Figures 1C, E and Figure 2, on the other hand, the network had only plastic synapses but no static synapses, i.e., *K* = 0. Without the static excitatory-inhibitory synaptic connections, as in Figure 1, the network lacked internally generated noise. Therefore, we injected external noise to the neuron’s voltage equation to induce variability in spiking activity (Fig. 2C). The variance of this white noise, *σ*^2^, was chosen such that the externally injected noise was similar to internally generated fluctuating synaptic current when static connections were present (*K* = 1000).

### 4.3 Network dynamics

In the following mathematical description of network activity, the static connections present in Figure 1B (*K* > 0) produced the balanced inputs 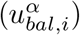, which exhibited large fluctuations. For this reason, no additional external noise was injected to the network (*σ* = 0). On the other hand, as mentioned above, in Figure 2 (and also in Figs. 1C, E), the static connections did not exist (*K* = 0) and external noise was injected to the neurons (*σ >* 0).

We used an integrate-and-fire neuron to model the membrane potential dynamics of *i*’th neuron:

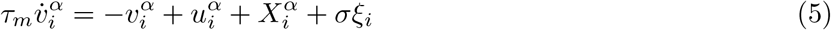

where a spike is emitted and the membrane potential is reset to *v*_*reset*_ when the membrane potential crosses the spike-threshold *v*_*thr*_. The membrane equations (Eq. 5), and the following synaptic equations (Eq. 8), were numerically solved using the forward Euler method.

Here, 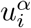 is the total synaptic input to neuron *i* in population *α* that can be divided into static and plastic inputs incoming through the static and plastic synapses, respectively:

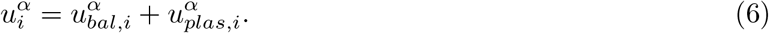

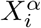 is the total external input that can be divided into constant external input, plastic external input, and the stimulus:

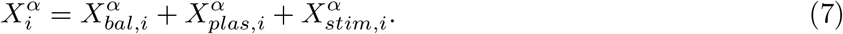

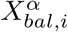 is a constant input associated with the initial balanced network. It scales with the number of connections (i.e. is proportional to 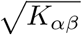), determines the firing rate of the initial network and stays unchanged [20].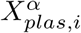 is plastic input provided to trained neurons in the recurrent network from external neurons that emit stochastic spikes with pre-determined rate patterns. The synaptic weights from the external neurons to the trained neurons were modified by the training algorithm. 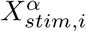 is the predetermined stimulus, generated independently from the Ornstein-Uhlenbeck process for each neuron, and injected to all neurons in the network to trigger the learned responses in the trained neurons.

The synaptic activity was modeled by an instantaneous jump of the synaptic input due to a presynaptic neuron’s spike, followed by an exponential decay. Our network did not include time delays in the propagation of the presynaptic neuron’s spike to the post-synaptic neuron; therefore a spike caused jumps in the postsynaptic neurons instantaneously. Since the static and plastic synapses did not overlap, we separated the total synaptic input into static and plastic components as mentioned above:

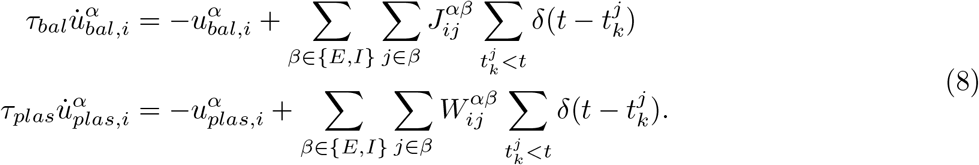

Alternatively, the synaptic activity can be expressed as a weighted sum of filtered spike trains because the synaptic variable equations (Eq. 8) are linear in **J** and **W**:

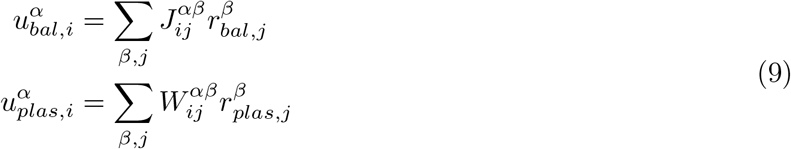

where

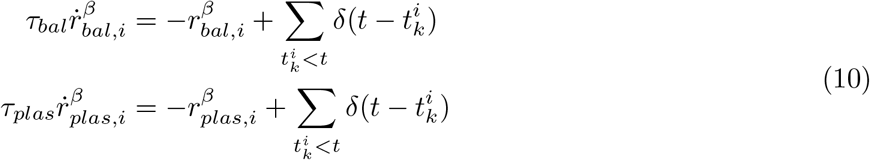

describe the dynamics of synaptically filtered spike trains.

Each external neuron emitted spikes stochastically at a pre-defined rate that changed over time. The rate patterns, followed by the external neurons, were randomly generated from an Ornstein-Uhlenbeck process with mean rate of 5 Hz. The synaptically filtered external spikes were weighted by plastic synapses **W**_**X**_ and injected to trained neurons:

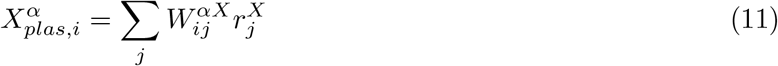

where

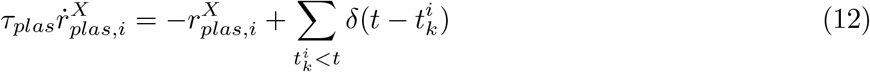

Similarly, the external stimulus *X*_*stim,i*_ applied to each neuron *i* in the network to trigger the learned response is generated independently from the Ornstein-Uhlenbeck process.

In the following section, we will use the linearity of **W, W**_**X**_ in Eqs. 9 and 11 to derive the training algorithm that modifies plastic synaptic weights.

### 4.4 Recursive least squares

Here we derive a synaptic update rule that modifies the plastic synapses to learn the target activities. The derivation closely follows previous papers [8, 16, 9]. For notational simplicity, we drop the neuron index *i* in **w**_*i*_ and other variables (e.g., *f*_*i*_, *u*_*i*_) but the same synaptic update rule is applied to all trained neurons. In fact, the plastic connections to every trained neuron can be updated simultaneously since each trained neuron has its own private target function and plastic connections. Therefore, the CPU multithreading (Fig. 1D) or GPU implementation (Figs. 1B, C) presented in our study leverages the fact that the synaptic update rule can be applied in parallel to all plastic synapses.

The cost function of each trained neuron *i* is defined by

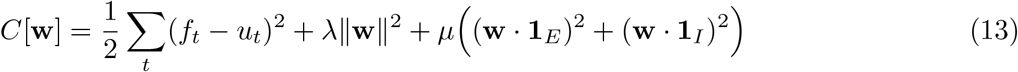

Specifically, 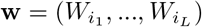 is a vector of recurrent plastic presynaptic synapses indexed by *i*_1_, …, *i*_*L*_; *f*_*t*_ is the target function over time *t*; *u*_*t*_ is the total synaptic input over time *t*. The hyperparameters *λ* and *µ* are the standard *L*_2_ and ROWSUM [16] regularization terms, respectively. The *L*_2_ regularization controls the overall learning rate, and ROWSUM regularization constrains the strength aggregate excitatory and inhibitory synaptic weights to the trained neuron.

The gradient of the cost function with respect to the weights **w** is

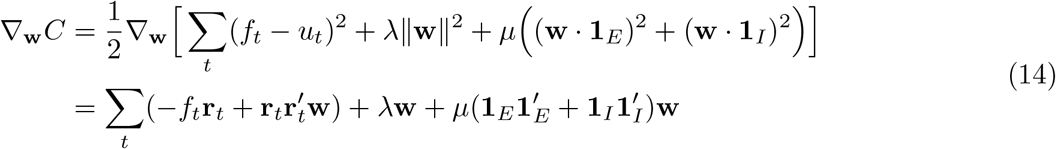

where we substitute the expression *u*_*t*_ = **w r**_*t*_ in the first line to evaluate the gradient with respect to **w**. To derive the synaptic update rule, we compute the gradient at two consecutive time points

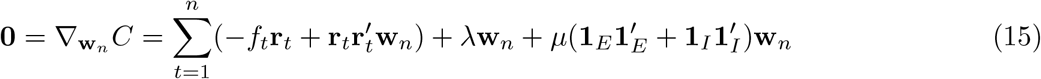

and

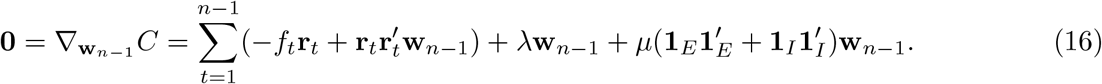

Subtracting Eqs (15) and (16) yields

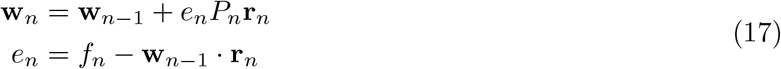

where

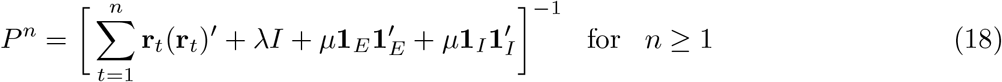

with the initial value

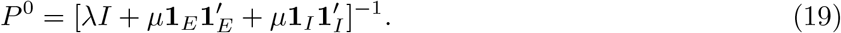

To update *P* ^*n*^ iteratively, we use the Woodbury matrix identity

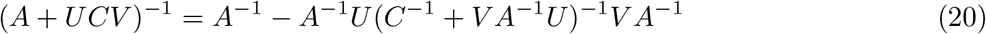

where *A* is invertible and *N* ×*N, U* is *N* ×*T, C* is invertible and *T* ×*T* and *V* is *T*× *N* matrices. Then *P* ^*n*^ can be calculated iteratively

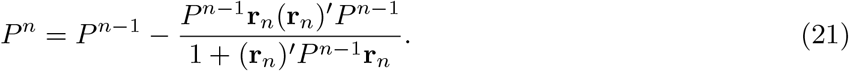

The following pseudocode provides overview of the simulatio function of the leaky integrate-and-fire neuronnand RLS training of spiking neural networks.

#### Algorithm 1

Simulation of spiking neural networks with RLS training

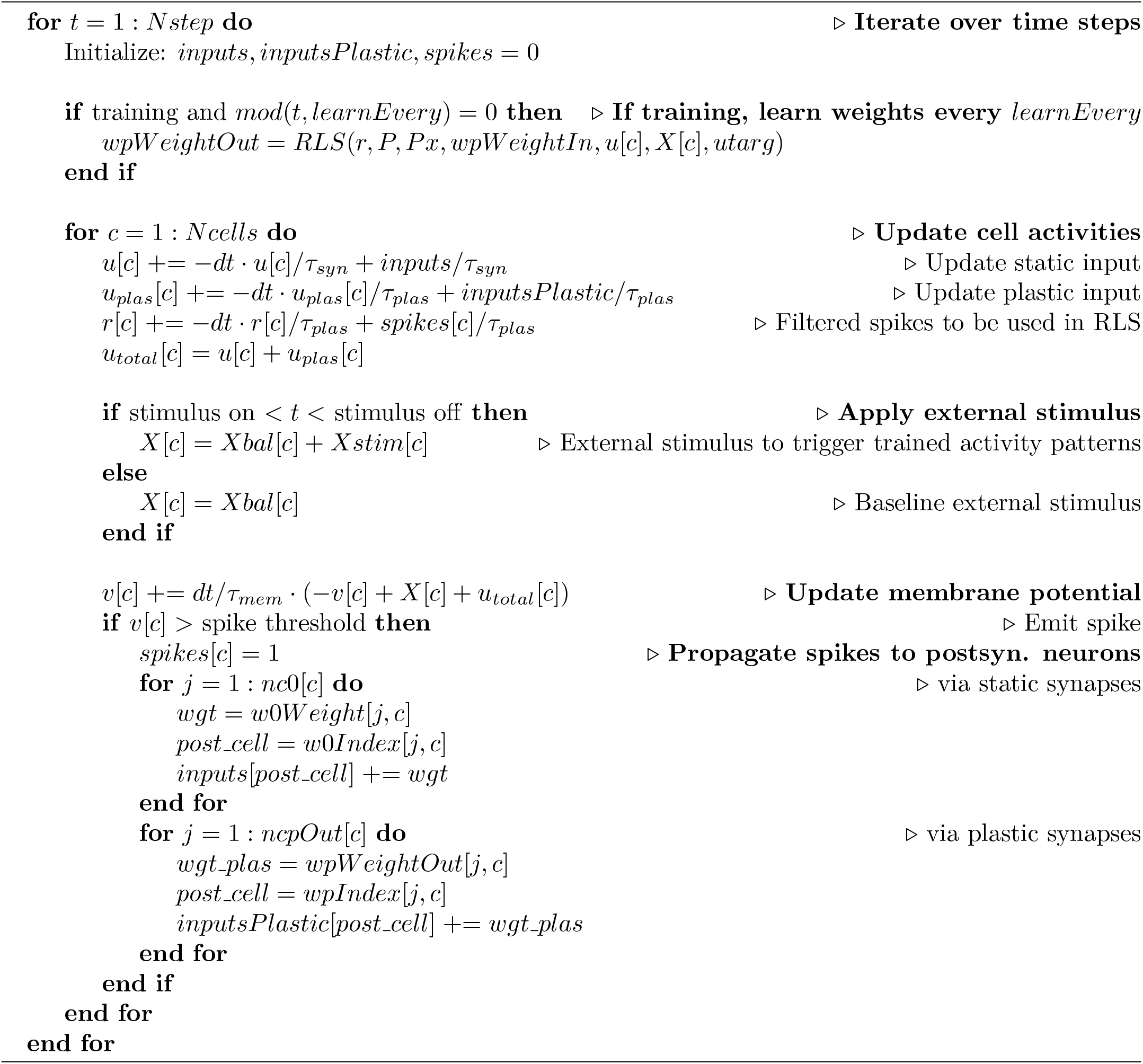

#### Algorithm 2

Recursive least squares algorithm

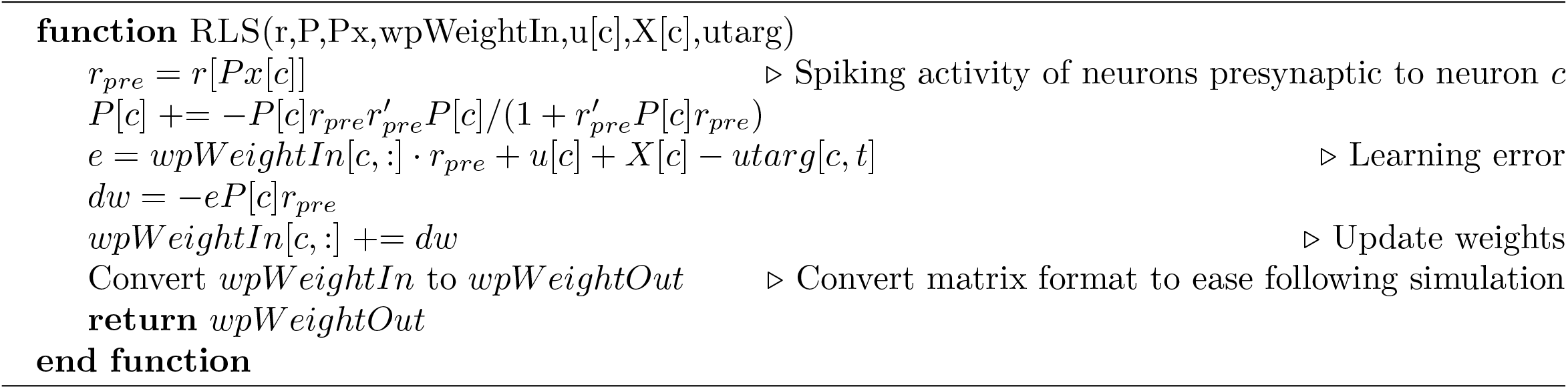

